# Copper chelation inhibits TGF-*β* pathways and suppresses epithelial-mesenchymal transition in cancer

**DOI:** 10.1101/2022.10.03.510707

**Authors:** E. M. Poursani, D. Mercatelli, P. Raninga, J. L. Bell, F. Saletta, F. V. Kohane, Y. Zheng, J. Rouaen, T. R. Jue, F. T. Michniewicz, E. Kasiou, M. Tsoli, G. Cirillo, S. Waters, T. Shai-Hee, E. Valli, M. Brettle, R. Whan, L. Vahadat, D. Ziegler, J. G. Lock, F. M. Giorgi, K. K. Khanna, O. Vittorio

## Abstract

Copper is a trace element essential to cellular function with elevated levels implicated in cancer progression. Clinical trials using copper chelators are associated with improved patient survival, however, the molecular mechanisms by which copper depletion inhibits tumor progression are poorly understood. This remains a major hurdle to the clinical translation of copper chelators. Epithelial-mesenchymal transition (EMT) is often exploited by malignant cells to promote growth and metastasis. Transforming growth factor (TGF)-*β* is a master regulator of EMT and facilitates cancer progression through changes in the tumor and its microenvironment. Herein, we report that a reduction of copper with the chelating agent tetraethylenepentamine (TEPA) inhibited EMT *in vitro* in three diverse cancer cell types; human triple-negative breast cancer (TNBC), neuroblastoma (NB), and diffuse intrinsic pontine glioma (DIPG) cell lines. Single-molecule imaging demonstrated EMT markers including Vimentin, *β*-catenin, ZEB1, and p-SMAD2 had increased expression with copper treatment and this pro-mesenchymal shift was rescued by the addition of TEPA. Moreover, SNAI1, ZEB1, and p-SMAD2 demonstrated increased accumulation in the cytoplasm after treating with TEPA. Transcriptomic analyses revealed a significant downregulation of the EMT pathway, including canonical (TGF-*β*/SMAD2&3) and non-canonical (TGF-*β*/PI3K/AKT and TGF-*β*/RAS/RAF/MEK/ERK) TGF signaling pathways. Matrix metalloproteinases MMP-9 and MMP-14 proteins which activate latent TGF-*β* complexes were also downregulated by TEPA treatment. These molecular changes are consistent with reduced plasma levels of TGF-*β* we observed in cancer models treated with TEPA. Importantly, copper chelation reduced metastasis to the lung in a TNBC orthotopic syngeneic mouse model. Our studies suggest copper chelation therapy can be used to inhibit EMT-induced metastasis by targeting TGF-*β* signalling. Because on-target anti-TGF-*β* therapies are failing in the clinic, copper chelation presents itself as a potential therapy for targeting TGF-*β* in cancer.

## Introduction

Copper is an essential catalytic cofactor involved in many biological processes in the human body (1, 2). Previous studies demonstrated that copper homeostasis is dysregulated in metastatic cancers (3). Copper accumulation has been observed in several malignancies such as breast, colon, gastric and lung cancer (4).

Copper chelation therapy was developed to treat Wilson's disease, a genetic disorder with an accumulative amount of copper (5). Copper chelators are approved for use in both adults and children and have recently attracted the attention of oncologists because of their proven ability to suppress tumor growth and angiogenesis in many pre-clinical studies (6, 7). We previously reported that copper chelation deregulated phosphorylation of STAT3 and AKT signaling which are important to the tumor and its microenvironment including blocking PD-1/PD-L1 axis in neuroblastoma (8, 9). Moreover, a recent study revealed that copper chelators inhibit tumor metastasis by collagen remodelling in the pre-metastatic niche in triple-negative breast cancer (TNBC) (7). However, the exact mechanism by which copper chelation reduces tumor metastasis is still poorly understood, with very few transcriptomic datasets published.

Epithelial-mesenchymal transition (EMT), is an embryonic phenotypic plasticity program which is reactivated in cancer cells in a dynamic fashion to acquire features that confer invasiveness, dissemination, and chemo/immunotherapy resistance (10, 11). EMT is the first and a critical step in developing metastasis and invasion in most types of human malignancies (12-14). Therefore, EMT represents an attractive target in cancer treatment. A number of important signalling pathways such as TGF-β, Wnt, Notch and Hedgehog are involved in modulating EMT via several transcription factors including smads, twist, snail and slug (15-17). Several studies have shown that TGF-β is a major regulator of the EMT process (18, 19). The canonical TGF-β signaling pathway acts through TGF-β binding its receptor, resulting in Smad2 and Smad3 (RSmads) phosphorylation at their C-terminal (20). Phosphorylation activates Smad2 and Smad3 to form a trimer with Smad4. The complex then translocates to the nucleus, where it regulates the transcription of target genes (19, 21). On the other hand, the non-canonical TGF-β signaling pathway stimulates phosphatidylinositol 3-kinase (PI3K) and its downstream pathways by phosphorylating AKT and promoting EMT signaling (14). Moreover, it was reported that TGF-β-induced cell migration mediated by MMP-14, which regulates MMP-9 activation (22, 23). Recent studies demonstrated a positive feedback loop between TGF-β and MMP-9 which is mediated by the PI3K signaling pathway (24). This confers acquired invasion and metastasis in cancer via induction of the EMT. TGF-β and MMP-9 are associated with migration and invasion, promoting metastasis in several malignancies (25). Moreover, there is an interconnection between TGF-β activation and MMP-9 expression, as an overexpression of MMP-9 increases the malignancy of breast cancer cell lines, largely via the activation of the SMAD signaling pathway (26).

TGF-β signaling has a central role in metastatic disease and is the subject of both molecular and clinical investigation over almost all cancer types (27). Breast cancer is the second-ranked malignancy leading to death among women worldwide (28, 29). Triple-Negative (ER-/PR-/HER2-) Breast Cancer (TNBC) is known to be the most aggressive among other subtypes of breast cancer with the least overall survival (30). TNBC is a heterogeneous tumor that remains clinically difficult to treat (31) and for which better therapeutic approaches are necessary. TGF-β acts as a driver of TNBC progression by inducing EMT and is involved in activating several MMPs, including MMP-9 and MMP-14 (13, 14). During breast cancer progression, TGF-β promotes metastasis by upregulating the expression of MMP-9 (32). Likewise, MMP-9, 2, and 14, can induce the biological activity of TGF-β via cleavage of latent TGF-β-binding protein-1 (33, 34).

Neuroblastoma (NB) is the most common extra-cranial solid tumor in children and accounts for 15% of childhood cancer deaths (35). Despite aggressive treatments, high-risk patients' survival rate is still poor (∼50%), and metastasis is one of the major causes of death in patients with NB (36). TGF-β signaling is involved in EMT in NB cells (37), and its upregulation caused increased migration and invasion *in vitro*. Moreover, MMP-9 has been recognized as essential factor in mediating drug and radio resistance and metastasis in NB patients (38).

Diffuse intrinsic pontine glioma (DIPG) is a lethal peadictric brain cancer with a median survial time ranging from 9 to 12 months, and has no curative treatments (39). On a molecular level, most DIPG have mutations in genes encoding Histone 3 mutations which often co-occurs with ACVR1 mutations. ACVR1 is part of the TGF-β superfamily of proteins and acts up stream of R-SMADs, suggesting DIPG’s have deregulated canonical TGF-β signaling. Moreover, DIPG cells have a mesenchymal, therapy-resistant phenotype which was found to be sensitive to chemical inhibition of AXL an initiator of the mesenchymal transition (40).

The tumor microenvironment of TNBC and NB is influenced by secreted TGF-β (41-43), making this cytokine a potential therapeutic target. Different strategies such as ligand traps, antisense oligonucleotides (ASOs), small molecule receptor kinase inhibitors, and peptide aptamers have been successfully used to target and modulate TGF-β signalling in cancers *in vitro*, however these strategies have faced severe problems in clinical trial studies (18, 44-46). 21 drugs targeting TGF-β have failed in clinical trials, partly because of severe side effects arising in patients (44-48).

In this study, we show for the first time that the copper chelator Tetraethylenepentamine (TEPA) significantly reduces TGF-β levels and subsequently inhibits both canonical and noncanonical TGF-β signaling pathways. TEPA downregulates proteins implicated in EMT-associated cancer invasion and metastasis in TNBC, NB and DIPG cell lines, and suppresses metastasis *in vivo*. Our data support the potential repurposing of copper chelators as a therapeutic strategy for targeting TGF-β in metastatic and invasive tumors.

## Materials & Methods

### Cell lines

The human triple-negative breast cancer cell line, MDA-MB-231, was kindly provided by Associate Professor Caroline Ford from UNSW (Sydney, Australia). Neuroblastoma cell line, SH-SY5Y (SY5Y) was sourced from ATCC with Loretta Lau. SH-SY5Y and MDA-MB-231 cells were cultured in DMEM, containing 2 mM L-glutamine (Gibco, Massachusetts, USA) and 10 % fetal bovine serum (FBS) (Sigma, Massachusetts, USA) and incubated in a humidified incubator with 5 % CO_2_ at 37 °C. HSJD-DIPG007 (DIPG007) cells were kindly provided by Dr. Angel Montero from the Hospital Sant Joan de Die in Spain. VUMC-DIPG010 (DIPG010) cells were kindly provided by Dr Esther Hulleman from the VU University Medical Centre in the Netherlands. Both DIPG lines were grown in a cancer stem cell media and growth factors (49) with 10% FCS to promote adherent cell growth.

### Reagents

Triethylenetetramine dihydrochloride (TEPA) was purchased from Sigma-Aldrich (Cat. No. T5033-25G). This powder is stable at room temperature. Stock solutions were prepared directly before use, as once in liquid it has an half-life of about 9 hours. Stock solution for *in vitro* work was prepared by dissolving the powder in complete cell culture media. For *in vivo* studies, TEPA was prepared in saline and administered by oral gavage.

### Immunofluorescence staining

MDA-MB-231 cells were seeded as 4000 cells/well in 96-well plate for an overnight and treated with 20 μM CuCL2, 15ng/mL TGF-β1, and 4mM TEPA in different conditions for 24 hours. Then, cells were fixed with 4% paraformaldehyde (Emgrid Australia, Cat# 15710) for 15 min and permeabilized with 0.1% Triton X-100 for 15 min at room temperature (RT). Following permeabilization, cells were blocked with Intercept (TBS) Blocking Buffer (LI-COR, Cat# LCR-927-60001) and incubated for 30 min at RT with the following primary antibodies: mouse anti-N-cadherin (1:100, Cell Signalling Technology, Cat# 14215S), mouse anti-vimentin (1:500, Abcam, Cat# ab8978), rabbit anti-SNAI1 (1:500, Proteintech, Cat# 13099-1-AP), rabbit anti-phosphorylated-AKT (Ser473) (1:500, Cell Signalling Technology, Cat# 3787), rabbit anti-β-catenin (1:500, ThermoFisher Scientific, Cat# 71-2700), rabbit anti-ZEB1 (1:200, Cell Signalling Technology, Cat# 3396) and rabbit anti-phosphorylated-SMAD2 (Ser465/467) (1:400, Cell Signalling Technology, Cat# 3108). Cells were then washed four times with Intercept Blocking Buffer, followed by a 15 min incubation with the secondary antibodies: anti-mouse AlexaFluor 488 (1:1000, Cell Signalling Technology, Cat# 4408) and anti-rabbit AlexaFluor 555 (1:1000, Cell Signalling Technology, Cat# 4413). Cell nuclei and actin filaments were stained simultaneously with 4′,6-Diamidino-2-phenylindole dihydrochloride (DAPI; 1:2000, Sigma-Aldrich, Cat# D9542) and Phalloidin-Atto 647N (1:2000, ATTO-TEC, Cat# AD647N) respectively. Before imaging, cells were washed four times with 1x TRIS buffered saline.

### Microscopy

Images were acquired using a Nikon AX R confocal microscope (Nikon, Tokyo, Japan) equipped with a 20x Plan Apochromat air objective (NA 0.75) at 2x zoom and resonant scanning at 2048×2048 resolution. 5 z-planes with 600 nm z-spacing were acquired per field and a maximum intensity projection was computed and used for subsequent image analysis steps. Channels were acquired in series with three passes: 405 nm and 647 nm in pass 1, 488 nm in pass 2 and 555 nm in pass 3.

### Image and data analysis

CellProfiler1 (Broad Institute, MIT, Massachusetts, USA) with Cellpose2 module integration was used for image analysis and quantitative feature extraction. Cell nuclei were detected using Cellpose model3 trained on DAPI images. DAPI and Phalloidin images were combined and then used to train another Cellpose model for the detection of cell bodies. The generated nuclei masks were subtracted from the cell body masks to produce cytoplasm masks. Mean total, nuclear, and cytoplasmic marker intensities were extracted for each cell.

Single cell data was parsed and analysed using custom-developed Knime workflows4 (KNIME AG, Zurich). Per-cell nuclear-to-cytoplasmic (N:C) ratios were calculated by dividing the mean nuclear intensity by the mean cytoplasmic intensity for each marker.

Outlier cells were detected and removed by means of interquartile range, R = [Q1 - k(IQR), Q3 + k(IQR)] with IQR = Q3 - Q1 and k = 1.5, using mean cell intensity and N:C ratio of each marker. Mean cell intensities were normalised via z-score. Pairwise Wilcoxon Rank Sum Tests were performed to compare fluorescence intensities between treatment groups.

### Cell Migration assay

Cell migration was assessed using the QCM Chemotaxis Cell Migration Colorimetric Assay kit (ECM506, Merck, USA). Briefly, the upper chamber is separated from a lower chamber with a microporous membrane (transwell insert) with a 5 μM pore size, which allows cells seeded in the upper chamber to migrate through the membrane towards lower chamber including chemoattractant, Briefly, MDA-MB-231 cells (1.5 × 10^5^ cells/well) were seeded in 6-well plate and incubated at 37°C for 4 hours. Then, MDA-MB-231 cells were treated with 0, 4, and 8mM TEPA for 24 hours prior to performing the migration assay. Then, the cells were harvested and seeded into the top chamber of the inserts containing low serum Media (0.5% FBS). The bottom chamber of each well included media with chemoattractant (20% FBS) along with corresponding TEPA concentrations that were consistent with the upper chambers. The migration plates were then incubated in a humidified incubator with 5 % CO_2_ at 37 °C for 16 hours. At the endpoint, transwell inserts were stained with crystal violet and imaged at 100x magnification using Nikon Eclipse microscopy (Molecular Devices, California, USA).

### Real-time monitoring of wound healing cell migration using IncuCyte imaging

MDA-MB-231 (2×10^4^ cells/well) and SH-SY5Y cells (4×10^4^ cells/well) were seeded in 96-well plates and cultured overnight to allow the cells reach confluency. The next day, an artificial gap was generated on a confluent monolayer of cells using an Incucyte® 96-well Woundmaker Tool (Satorious, Germany) and cells were washed twice with PBS. Cells were then treated with specific amounts of TEPA (0, 4, and 8 mM for MDA-MB-231, and 0 and 2 mM for SH-SY5Y). For the co-treatment experiment, TGF-β was used at 15 ng/mL for MDA-MB-231, and at 10 ng/mL for SH-SY5Y cells. After treatment plates were immediately incubated at 37°C, 5 % CO_2_. Cells were imaged every 2-4 hours. Images and data were obtained using IncuCyteS3 (Essen Bioscience) with a 10x object. Incucyte® Scratch Wound Analysis Software Module (Cat. No. 9600-0012) was used to automatically measure the scratch length.

### RNA sequencing

MDA-MB-231 human TNBC cells were plated at 5×10^6^ cells/T75 flask in DMEM media supplemented with 10% FBS and treated with 4 mM TEPA for 8 and 24 hours. DIPG010 cells were seeded in 6 well plates at 5×10^5^ cells and treated with 4 mM TEPA for 24 hours. Cells were then harvested, washed with PBS, and lysed using the Qiagen RNA Extraction Kit (RNEasy kit, catalog no. 74104, Qiagen, Hilden, Germany) following the recommended protocol. RNA purity and concentration were assessed using the Nanodrop 2000 (Thermo Fisher Scientific, USA). Quality control and library preparation were carried out by Ramaciotti Centre for Genomics (UNSW, Australia). Paired-end 100-bp sequencing was performed on the Illumina NovaSeq 6000 using TruSeq stranded mRNA. The data have been deposited under Gene Expression Omnibus (GEO) accession number GSE185760. We also analyzed our RNA-seq data obtained from SH-SY5Y treated with TEPA (accession number: GSE155031).

### Bioinformatics analysis

After a quality check performed with FastQC (50), paired-end RNA-seq data were aligned to the human genome assembly (build hg38) with the HISAT2 alignment program (51). Raw gene counts were obtained by running featureCounts (52) with the following settings: -p -s 2 -t exon -g gene_name. Reads were mapped to genomic features using Homo_sapiens.GRCh38.103.gtf (53). Differential expression analysis was performed using DESeq2 (v1.32.0) (54) in R (55). Enrichment analysis was performed using the enrichR package (v3.0) (56), while Gene Set Enrichment Analysis (GSEA) was performed using fgsea (v1.18.10) R package and the human gene sets from the Broad Institute's MSigDB, accessed through the msigdbr package (v7.4.1) (57-59). Enrichment plots were generated using plotting functions from the corto package (1.1.8) (60). The Gene expression matrix was obtained by applying Variance Stabilizing Transformation (VST) to raw counts data before plotting (61). SH-SY5Y expression matrix was obtained from GEO (accession number: GSE155031) (8).

### Western blot analysis

MDA-MB-231 and SH-SY5Y cell lines were seeded in 6-well plates (2×10^5^ cells/well) overnight and then were treated with specific amounts of TEPA (0, 4, and 8 mM for MDA-MB-231, and 0 and 2 mM for SH-SY5Y). After 24 and 48 hours, cells were harvested and lysed in cold RIPA lysis buffer supplemented with 1X cOmplete Mini, EDTA-free Protease Inhibitor Cocktail, and PhosSTOP™ phosphatase inhibitor (both Sigma-Aldrich, Schnelldorf, Germany). DIPG cell lines were seeded in 6-well plates (5×10^5^ cells/well) overnight and then were treated TEPA. Protein content was measured using BCA Protein Assay Kit (Thermo Fisher Scientific, USA). An equal amount of protein from control and each experiment was loaded onto 4-20% SDS-PAGE gels (Mini-PROTEAN TGX Gels, Bio-Rad, USA) and transferred to the nitrocellulose membranes (Amersham™ Protran® 0.45 μM, GE Healthcare, Germany). Nitrocellulose membranes containing separated proteins were blocked in 5% BSA/TBS-T (1X TBS + 0.1% Tween-20) for 1 hour and probed with antibodies against Smad2 (ab40855, Abcam), phospho-Smad2(Ser465/467) (138D4, Cell Signaling Technology), AKT (pan) (11E7, Cell Signaling Technology), phospho-AKT(S473) (193H12, Cell Signaling Technology), Phospho-p44/42 MAPK (pERK1/2)(T202/Y204) (D13.14.4E, Cell Signaling Technology), p44/42 MAPK (ERK1/2) (9102S, Cell Signaling Technology), MMP-9 (D6O3H, Cell Signaling Technology), MMP-2 (A6247, ABclonal), MMP-14 (A2549, ABclonal), mTOR (2972, Cell Signaling Technology), Phospho-mTOR(Ser2448) (2971, Cell Signaling Technology), E-cadherin (24E10, Cell Signaling Technology), Vimentin (D21H3, Cell Signaling Technology), and GAPDH (sc-365062, Santa Cruz) overnight at 4°C. After washing in TBS-T, membranes were incubated for 1hr at room temperature with polyclonal HRP-conjugated goat anti-rabbit (1:5000; Dako, Denmark) or anti-mouse (1:5000; Glostrup, Denmark) IgG secondary antibodies. The resultant immunoreactive band signal was visualized using Clarity ™ Western ECL Substrate (Bio-Rad, USA). The accurate quantification of the relative density between samples was analysed by Image Lab™ software v6.0 (Bio-Rad, California, USA) and normalized with GAPDH, as a housekeeping protein.

### RNA extraction & quantitative PCR (qPCR)

MDA-MB-231, DIPG and SH-SY5Y cell lines were seeded and treated as previously described in the western blot section. Total RNA was extracted using the RNeasy Mini Kit (#74106; QIAGEN, California, USA) according to the manufacturer's instructions and its quality and concentration was assessed using the Nanodrop 2000 (Thermo Fisher Scientific, USA). First strand cDNA was synthesised with 1μg of total RNA using SuperScript II Reverse Transcriptase (Invitrogen, Massachusetts, USA) according to the manufacturer’s instructions. 40 ng of the resulting cDNA was used in the subsequent qPCR analysis using SsoAdvanced Universal SYBR® Green Supermix (#1725270) and CFX96 Real-Time PCR Detection System (both Bio-Rad, USA) with the following PCR program: 95°C for 3 min, followed by 40 cycles of 95°C for 10 s, 60°C for 30 s and 72°C for 30 s). Primer sequences were as follows: *MT1X* (F: GCTTCTCCTTGCCTCGAAAT and R: GCAGCAGCTCTTCTTGCAG), *TGF-β* (F: AAGTGGACATCAACGGGTTC and R: GTCCTTGCGGAAGTCAATGT) and *GUSB* (F: TGGTGCGTAGGGACAAGAAC and R: CCAAGGATTTGGTGTGAGCG). Quantifications were normalized using *GUSB* as the endogenous control.

### *In vivo* xenografts and tumor growth analysis

All experiments were approved by the QIMR Berghofer Medical Research Institute Animal Ethics Committee and performed as per the committee’s guidelines (62). For murine 4T1.2 and 4T1BR4 TNBC syngeneic models, 1×10^5^ cells were prepared in PBS and injected into the right 4^th^ inguinal mammary fat pad of 6-weeks old female immunocompetent Balb/c mice. Once tumor size reached 50 mm^3^, mice were randomized blindly into different treatment groups and were then treated with the vehicle (saline) or TEPA. Two treatment groups were created, one was treated with 400 mg/kg of TEPA and the second with 800 mg/kg of TEPA as daily, oral gavage which was continued for three weeks. Tumor growth was measured thrice weekly using a digital caliper. Tumor volume was measured using the following formula: tumor volume = [L×W^2^]/2, where W = width of the tumor and L = length of the tumor.

### *In vivo* lung metastases analysis

4T1BR cells were injected into 6-weeks old female Balb/c mice as described above. Once tumor size reached ∼30-50 mm^3^, mice were randomized blindly into different treatment groups and were then treated with the vehicle (saline) or TEPA (400 mg/kg, oral gavage, daily) for three weeks. This concentration was used as it did not impact the primary tumor growth in this model. At the end of the treatment, all mice were euthanized, and their lungs were harvested. For lung metastasis, lungs were perfused with 1X PBS to remove residual blood; then they were fixed with 10% formalin and stained with hematoxylin and eosin (H&E) to facilitate the counting of micro- and macro-metastatic nodules.

### Measurements of TGF-β in mice sera

The detection of TGF-β in the cell culture media from SH-SY5Y and MDA-MB-231 cells treated with TEPA for 24h was performed according to the protocol suggested by Thermo Fisher (TGF-beta-1-Human-ELISA-Kit/BMS249-4).

For the *in vivo* experiments in the 4T1.2 syngeneic model, whole blood was obtained from the tail vein of mice 30 days after commencing treatment with TEPA 800mg/Kg. Serum was obtained through centrifugation (15min at 2500 RPM) of clotted blood and stored at –70°C until use. TGF-β levels (pg/mL) were determined using a commercial murine ELISA kit (#BMA608-4; Invitrogen, Massachusetts, USA) according to the manufacturer’s instructions. Samples were assayed in duplicate and expressed as the average reading using a Benchmark Plus microplate spectrophotometer reader (Bio-Rad, California, USA) at 450 nm (620 nm reference wavelength).

### Statistical analysis

Statistically significant differences were determined if the p-value by an ordinary one-way ANOVA and paired t-test was less than or equal to 0.05 (Prism version 8.3.1).

## Results

### Copper homeostasis affects expression and cellular localization of EMT markers

Copper-chelation in preclinical models of cancer has been shown to suppress metastasis in-part through impact on tumor stroma (63). However, tumor intrinsic effects of copper-depleted metastatic propensity and changes in the molecular markers of EMT in cancer cell lines have not been investigated. To investigate whether intracellular copper levels alter EMT markers we treated MDA-MB-231 cells with copper and used single molecule confocal microscopy to analyse cells (Fig1). Mesenchymal protein markers vimentin, β-Catenin, ZEB1, pSMAD2, and N-cadherin were significantly increased by copper treatment compared to untreated cells. This effect was completely rescued by the addition of the copper chelator TEPA. TEPA treatment on its own had a significant decrease of the above protein markers compared to control, suggesting endogenous intracellular copper could be further targeted to reduce mesenchymal phenotype. Unlike the other factors, SNAI1 protein did not increase significantly with copper treatment, but did reduce with TEPA treatment. Perhaps this is because these cells already are highly mesenchymal-like and SNAI1 protein endogenous levels are close to their maximum and cannot easily be increased. Importantly, for SNAI1, ZEB1 and pSMAD2, confocal microscopy showed not only that the total level of these proteins decreased with TEPA treatment but that the nuclear:cytoplasmic ratio of their expression was more cytoplasmic after treatment. As a positive control for mesenchymal modulation of cell type, we also treated cells with recombinant TGF-β. In accordance with TGF-β’s role in the promotion of EMT, all the mesenchymal markers analysed displayed an increase in expression after TGF β treatment.

**Figure 1.**
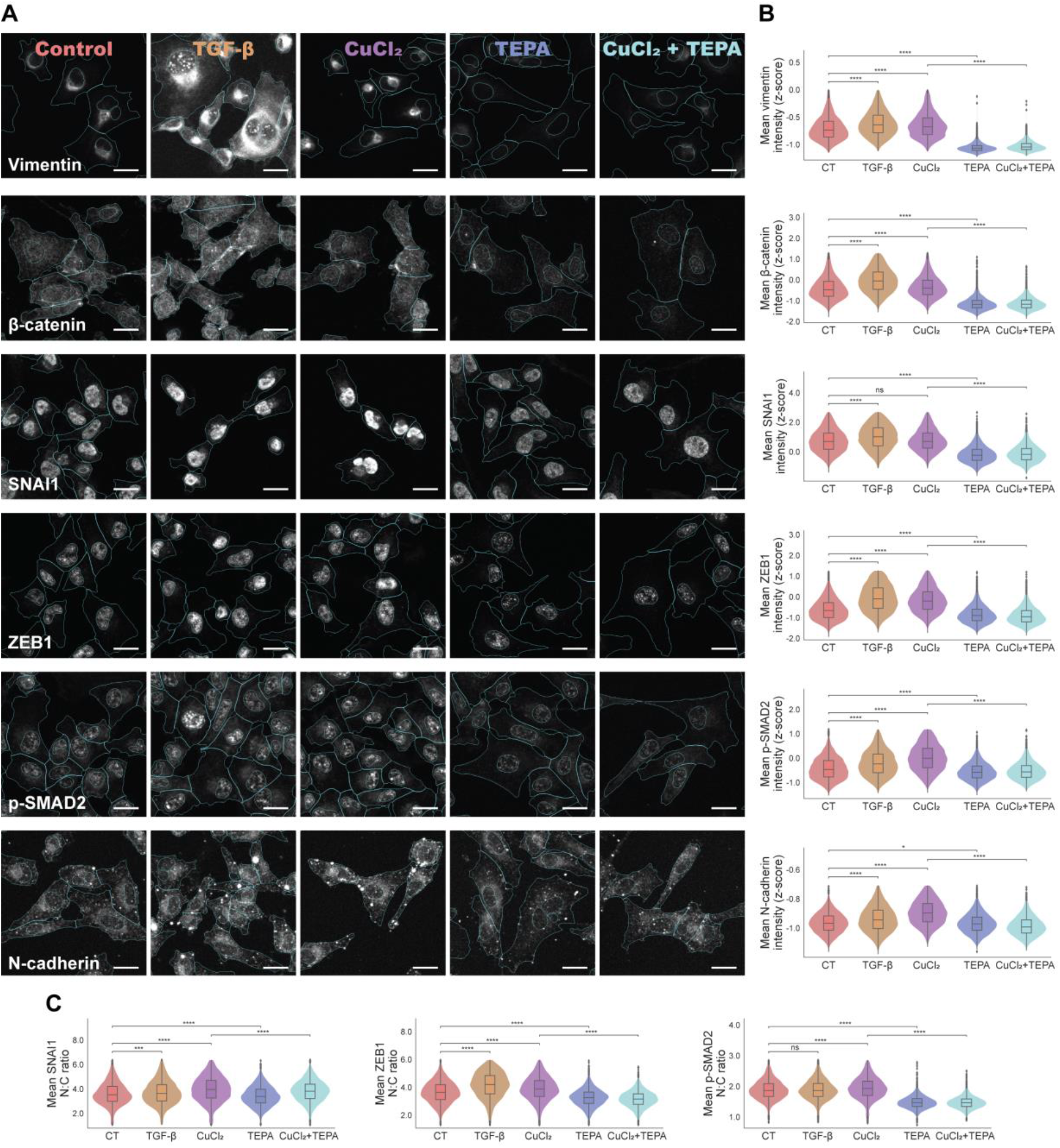
a Expression of various EMT markers in MDA-MB-231 TNBC cells treated with TGF-β, CuCl2, TEPA and CuCl2+TEPA. Cell nuclei and cytoplasm outline shown in cyan. All scale bars 20 μm. b Quantification of single cell mean fluorescence intensities. C: Quantification of single cell mean nuclear-to-cytoplasmic (N:C) ratio. Total cells per treatment condition: control = 21848, TGF-β = 17743, CuCl2 = 19153, TEPA = 25200, and CuCl2+TEPA = 18705. 14000-23000 cells per marker, 2000-6000 cells per marker per treatment group. Two-sided Mann-Whitney-Wilcoxon test with Bonferroni correction, ns: (5.00e-02 < p <= 1.00e+00), *: (1.00e-02 < p <= 5.00e-02), **: (1.00e-03 < p <= 1.00e-02), ***: (1.00e-04 < p <= 1.00e-03), ****: (p <= 1.00e-04).

### TEPA inhibits cell migration and invasion in TNBC and NB cell lines

To investigate whether TEPA inhibits EMT-induced migration and invasion in cancer cells, we first studied the effect of TEPA on proliferation, which was not impacted after treating cells with increasing concentration of TEPA (Suppl. Fig. 1a). We next performed migration assay for MDA-MB-231 TNBC cells by pre-treating them different doses of TEPA. Cell migration was significantly inhibited by TEPA in a dose-dependent manner in TNBC cells (Fig.2a). Treatment with 4 and 8mM of TEPA inhibited MDA-MB-231 cell migration by 21% and 45%, compared to their respective non-treated control cells respectively. This result was strengthened by additional data obtained from the transwell cell invasion assay showing the ability of TEPA to inhibit the invasion of cancer cells through the extracellular matrix, a critical process for metastasis initiation (Fig. 2b). Moreover, we found that chelating copper with TEPA alters the morphology of cells morphology and increased the area of the cells (Fig. 2c). To further prove that TEPA reduced cancer cells migration, we performed a scratch-wound healing assay in both MDA-MB-231 TNBC and SH-SY5Y NB cell lines. In Suppl Fig. 1b and 1c demonstrate that TEPA inhibited cell migration at non-toxic concentrations in both cell lines. Moreover, to understand if supplementation of TGF-β could accelerate cancer cell migration and if TEPA was able to counteract its effect, TGF-β was added to the cell culture media to accelerate the wound healing process in a scratch assay. It is important to highlight that TEPA maintained an inhibitory effect on cell migration in both TNBC and NB tumor cell lines, even after stimulation with TGF-β (Fig. 2d & 2e).

**Figure 2.**
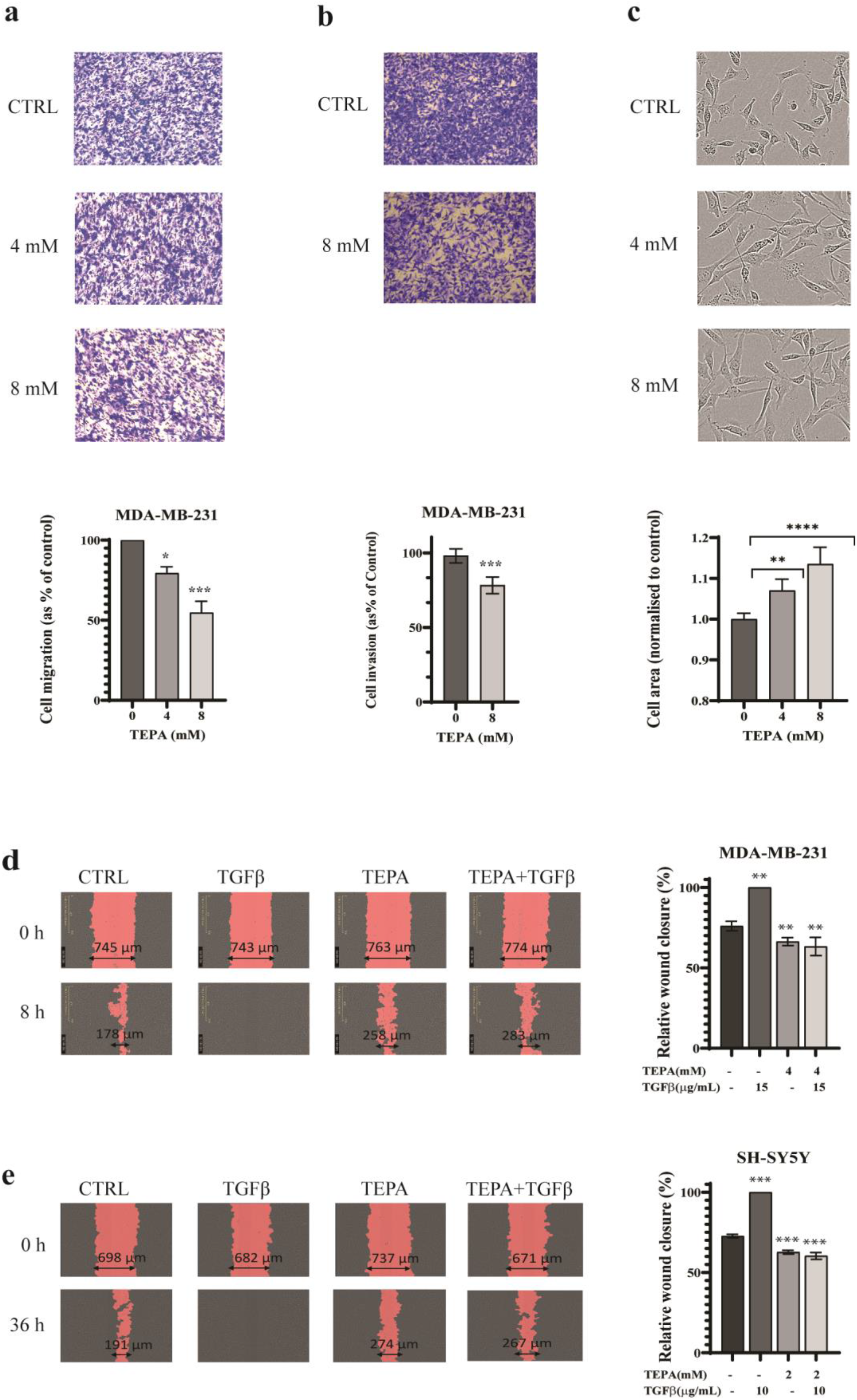
Inhibitory effect of TEPA on cell migration and morphology in TNBC (MDA-MB-231) and NB (SH-SY5Y) cells. **a** Representative images (n=3) of transwell migration assays performed on MDA-MB-231 cells treated with 0, 4, and 8 mM of TEPA. **b** Transwell cell invasion assay for MDA-MB-231 cells treated with 0 and 8 mM of TEPA. **c** Representative morphology alteration of MDA-MB-231 cells treated with TEPA. Images show how TEPA treatment caused a transition from a mesenchymal to an epithelial-like phenotype. Images were taken at 100x magnification using an ECLIPSE Ni-E light microscope. The graph shows the average cell size (μm2) relative to the control. Significance was determined by One-way ANOVA where ** and **** represent p<0.001 and p<0.0001, respectively. **d & e** Inhibitory effect of TEPA on cell migration induced by TGF-*β* in a scratch assay. After generating artificial gaps, TNBC and NB cells were pre-treated with TEPA (4mM for TNBC and 2mM for NB) for two hours and then with 15 ng/ml TGF-*β* for MDA-MB-231 and 10 ng/ml TGF-*β* for SH-SY5Y in complete media. Wound closure was monitored by IncuCyteS3 at 0 and 14 hours for TNBC and 0 and 36 hours for SH-SY5Y. Significance was determined by One-way ANOVA with p-value <0.0001 for both MDA-MB-231 and SH-SY5Y cells.

### TEPA downregulates the EMT pathways in cancer cell lines

Copper chelation has never been shown to alter the mesenchymal phenotype, thus we chose to map the underlying molecular mechanisms by which TEPA inhibits EMT in diverse cancer types. We performed whole transcriptome analysis on TEPA treated TNBC, NB, and DIPG cells (Fig. 3, 4). Our fundings confirmed that TEPA treatment reverses the mesenchymal-like phenotype of the TNBC cells (Fig. 3) and DIPG cells (Fig. 4). Differential expression analysis comparing TEPA treated cells to untreated controls revealed that TEPA treatment resulted in a robust gene expression signature, significantly affecting the expression of a total of 2649 genes at an early (8 hours), and a total of 2823 genes at a late (24 hours) time points in MDA-MB-231 cells (Suppl Fig. 2a & 2b). Unsupervized clustering grouped differentially expressed genes into four clusters, based on their time-dependent response to TEPA at different treatment time-points (T0 = 0h, T1 = 8h; T2 = 24 h): 1180 genes show a tendency to downregulation (cluster 1), 1248 genes are temporarily upregulated before returning essentially to pre-treatment levels (cluster 2), 820 genes are temporarily downregulated before returning to pre-treatment levels (cluster 3), and 1242 genes are continuously upregulated (cluster 4) by TEPA treatment (Suppl Fig. 2c). We further investigated the effect of TEPA on molecular pathways by performing a global Gene Set Enrichment Analysis (GSEA). Our analysis revealed a consistent and significant deregulation of the Hallmark Epithelial to Mesenchymal Transition pathway after TEPA treatment, both at 8 and 24 hours in MDA-MB-231, including the downregulation of MMP2, MMP9, MMP14, PLOD2, PLAT, MARCKSL1, CCR7, PAK1, ZYX, and an upregulation of E-cadherin (CDH1), TIMP1 and TIMP3 (Fig. 3a & 3b). Similar to the observation from the TNBC transcriptomic analyses, 24 hours of TEPA treatment in a DIPG brain cancer patients derived cell line, DIPG010 showed a downregulation of mesenchymal associated transcripts including Vimentin (VIM), MMP2, N-cadherin (CDH2), VCAN, PLOD2, ZYX and PLAT (Fig. 4a). In the TNBC cells, expression of the leading-edge genes, i.e. the subset of genes mostly contributing to the enrichment of transcripts in the EMT pathway, are shown in the heatmap in Fig. 3a and similarly in Fig. 3b and Fig. 4a. Among others, downregulation of lysyl oxidases, metallopeptidases and several crucial components of TGF-β signaling were detected in MDA-MB-231 (e.g., downregulation of TGFBR2, SNAI2, SPARC, RAC2, and BCL2, and upregulation of CDKN1A; Table 1) as well as in DIPG010 (e.g., downregulation of SPARC, Vimentin, and TGF-β1). Interestingly, a time-dependent downregulation trend was detected for SNAI2 (Snail Family Transcriptional Repressor 2, also known as SLUG). This is a well-studied transcriptional regulator of EMT its inhibition has been previously shown to decrease TNBC dissemination in animal models (64, 65). Downregulation of SNAI2 at gene network level was confirmed by Master Regulator Analysis at both time points (Suppl. Fig. 3). The Master Regulator Analysis algorithm implemented in corto (60) allows the analysis of gene regulatory networks to identify master transcriptional factors that control specific groups of genes explaining an observed phenotype (66). Importantly, we detected a significant downregulation of the VERRECHIA EARLY RESPONSE TO TGF-β1 pathway at both time points, indicating inhibition of TGF-β signaling upon TEPA treatment in TNBC cell lines (Fig. 3c & 3d). We investigated whether the effects of TEPA on TNBC cells’ transcriptome affecting the pathways discussed above could represent a general mechanism of action for TEPA, or if those changes would represent a restricted mechanism observed in TNBC cells.

**Table 1.** Gene signature of several important cancer-related signaling pathways in MDA-MB-231 TNBC cells treated with specific doses of TEPA for 8 and 24 hours. Genes with blue color are downregulated, and genes with red color are upregulated.

| Signalling pathways (KEGG) | Genes Symbol |  |
| --- | --- | --- |
|  | 8 hours | 24 hours |
| TGF-β/SMAD2/3 pathway | TGFBR2, SPARC | SNAI2, RAC2, BCL2, **CDKN1A** |
| PI3K/AKT pathway | PIK3C2B, PIK3IP1, HSP90 |  |
| mTOR pathway | TNFRSF1A | WNT5A, eIF4E |
| MAPK pathway | TGF-α, MKNK2, DUSP6, TNFRSF1A | RAC2 |
| Ras/Raf/Erk signalling pathway | TIAM1 | RASSF5, IKBKG |
| TLR/NFκB pathway | TLR2, TLR3, TRAF6, RAC2, MAP2K6, RIPK1 | IKBKG, PLAU |
| IGF signalling pathway | IGFBP-1, IGFBP-4, IGFBP-5, IGFBP-6, IGFBP7 | IGFBP-3, ITGA2, ITGA3 |
| ECM receptor signalling | ITGA1, ITGAV, ITGB3, ITGB5 | COL4A1 |
| Focal adhesion |  | PXN, LAMA4, ZYX |
| Inflammation & immune system suppression | PTX3, CXCL1, CXCL8, IL15, IL6 | IL6R |
| Cell mobility, invasion & metastasis | PLOD2, PLAT, MARCKSL1, CCR7, MMP11, MMP15 | PAK1, ZYX, **TIMP3**, **TIMP4**, **CDH1** |
| Apoptosis and cell cycle inhibition | BCL2L1 | BCL2, **CDKN1A**, **CDKN2D** |
| WNT signalling | NOTCH3, WNT9A, FZD1, FZD8, AXIN2, PFN2 | NOTCH1, WNT5A, WNT7B, WNT10B |
| Angiogenesis | VEGFB |  |

**Figure 3.**
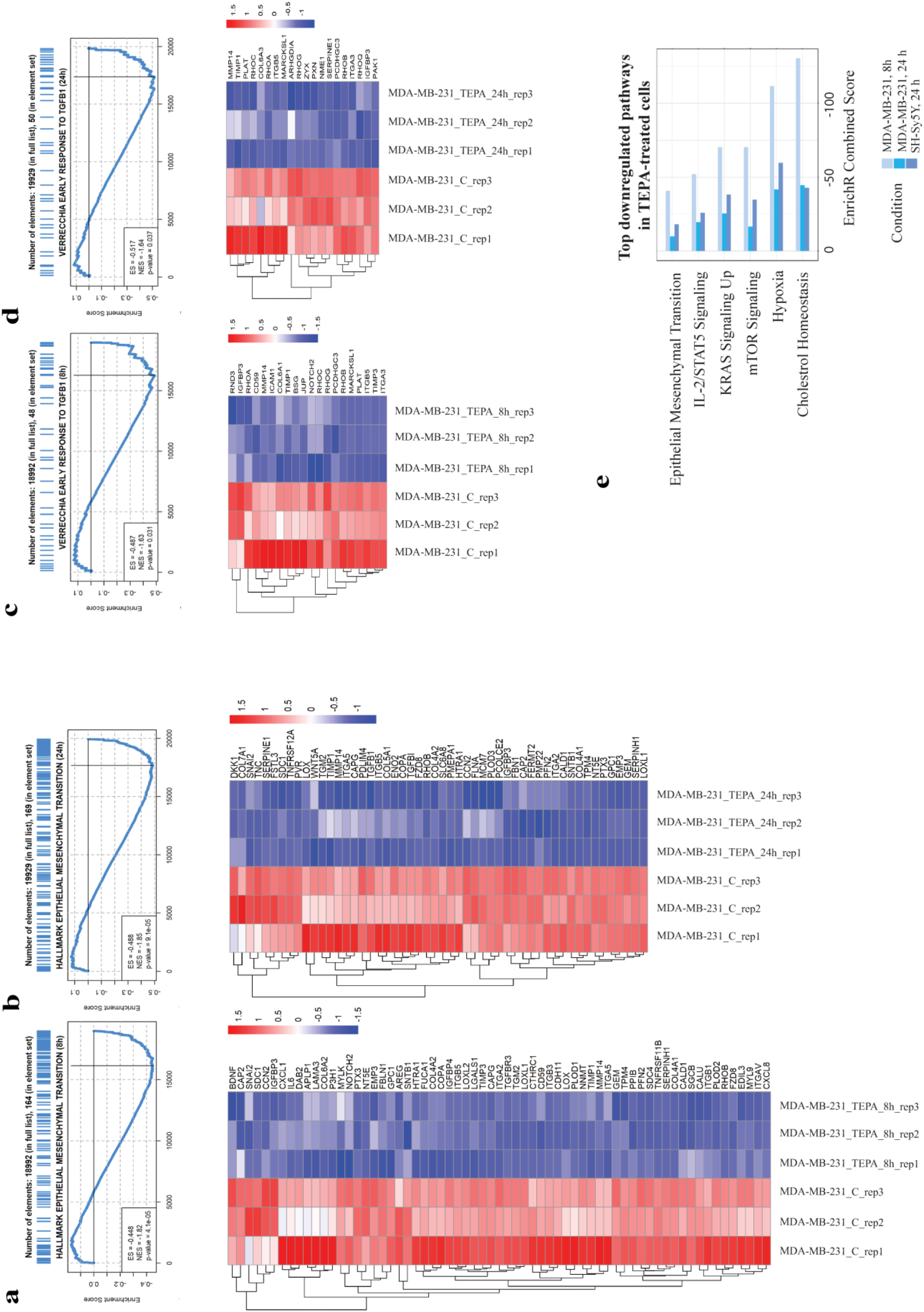
Enrichment analysis of TEPA-treated cells at 8 and 24 hours. a & b Up, GSEA enrichment plot of EMT-related genes identified by RNAseq analysis in MDA-MB-231 cells treated with TEPA for 8 (left panel) and 24 (right panel) hours. Downregulation of the MSigDB Hallmark EMT gene set is consistently observed at both time-points. Down, Heatmap showing EMT gene set leading-edge genes at MDA-MB-231 cells treated with TEPA for 8 (left) and 24 (right) hours. Heatmap colour intensity is proportional to row-scaled VST-normalized expression. c & d Up, GSEA enrichment plot showing downregulation of early TGFB1 early responsive genes identified by RNAseq analysis in MDA-MB-231 cells treated with TEPA for 8 (left panel) and 24 (right panel) hours. Down, Heatmap showing MSigDB’s VERRECCHIA EARLY RESPONSE TO TGFB1 gene set leading-edge genes at MDA-MB-231 cells treated with TEPA for 8 (left) and 24 (right) hours. The heatmap color intensity is proportional to row-scaled VST-normalized expression. e Barplot showing common negatively enriched gene terms in MDA-MB-231 cells treated with TEPA for 8 and 24 hours and SH-SY5Y treated with TEPA for 24 hours. Enrichment analysis was performed with the enrich R package. A consistent downregulation of EMT and cholesterol metabolism can be observed in both breast cancer and neuroblastoma cells. Furthermore, downregulation of several signaling pathways (IL-2/STAT5, mTORC1 and KRAS) is observed in both cell models following TEPA treatment.

**Figure 4.**
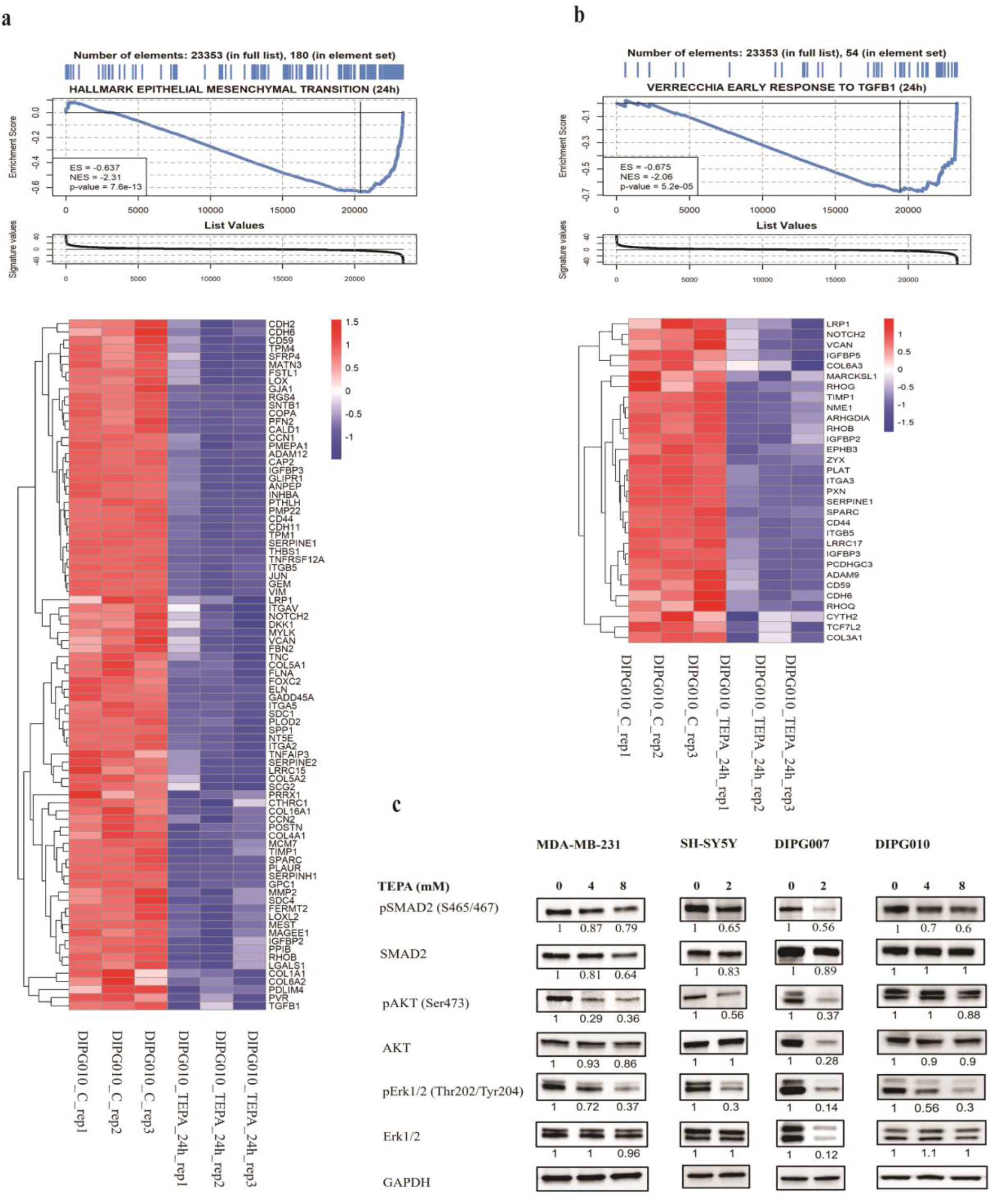
Enrichment analysis of TEPA-treated DIPG010 cells at 24 hours. a Up, GSEA enrichment plot of EMT-related genes identified by RNAseq analysis in DIPG010 cells treated with TEPA for 24 hours. Downregulation of the MSigDB Hallmark EMT gene set is consistently observed at this time-point. Down, Heatmap showing EMT gene set leading-edge genes at DIPG010 cells treated with TEPA for 24 hours. Heatmap colour intensity is proportional to row-scaled VST-normalized expression. b Up, GSEA enrichment plot showing downregulation of early TGFB1 early responsive genes identified by RNAseq analysis in DIPG010 cells treated with TEPA for 24 hours. Down, Heatmap showing MSigDB’s VERRECCHIA EARLY RESPONSE TO TGFB1 gene set leading-edge genes at DIPG010 cells treated with TEPA. The heatmap color intensity is proportional to row-scaled VST-normalized expression. c Analysis of TGF-*β*/SMAD2/3 and TGF-*β*/AKT/mTOR signaling pathways at protein levels. Western blot analysis of SMAD2, phospho-SMAD2, AKT, phospho-AKT, ERK1/2, and phosphor-ERK1/2 following treatment of MDA-MB-231, SH-SY5Y, DIPG007, and DIPG010 cells with TEPA (0, 4 & 8mM for MDA-MB-231 and DIPG010, and 0 & 2mM for SH-SY5Y and DIPG007) for 24 hours. GAPDH has been used as a positive and an endogenous control for normalization.

Our data reveal common negatively enriched gene terms in MDA-MB-231 and SH-SY5Y, and DIPG cells treated with TEPA. As Fig. 3e showed, a significant negative enrichment of EMT, hypoxia, cholesterol homeostasis, and IL-2/STAT5 pathways this suggests a downregulation of these processes in both cell types.

### TEPA blocks TGF-β canonical and noncanonical signaling pathways in cancer cell lines

TGF-β is a cytokine that acts on protein kinase receptors to induce a plethora of biological signals regulating EMT, angiogenesis, inflammation, and immune response. To explore potential crosstalk of the copper chelator with TGF-β signaling, we interrogated our RNAseq obtained from MB-231-MDA cells treated with 4mM TEPA for 8, 24 hours and SH-SY5Y cells with 2mM TEPA for 24 hours. TGF-β/SMAD and TGF-B/AKT/mTOR signaling pathways are displayed as deregulation in MB-231-MDA after 8 hours and 24 hours (Fig. 5a & 5b). Correlating experimental results with RNAseq data, we summarized in Fig. 5d the effect of TEPA on multiple factors causing the inhibition of TGF-β and its network of signaling pathways. In table 1, we have clarified the signaling pathways and their specific genes, which are downregulated by TEPA in TNBC cancer cells, 8- and 24-hours after treatment.

**Figure 5.**
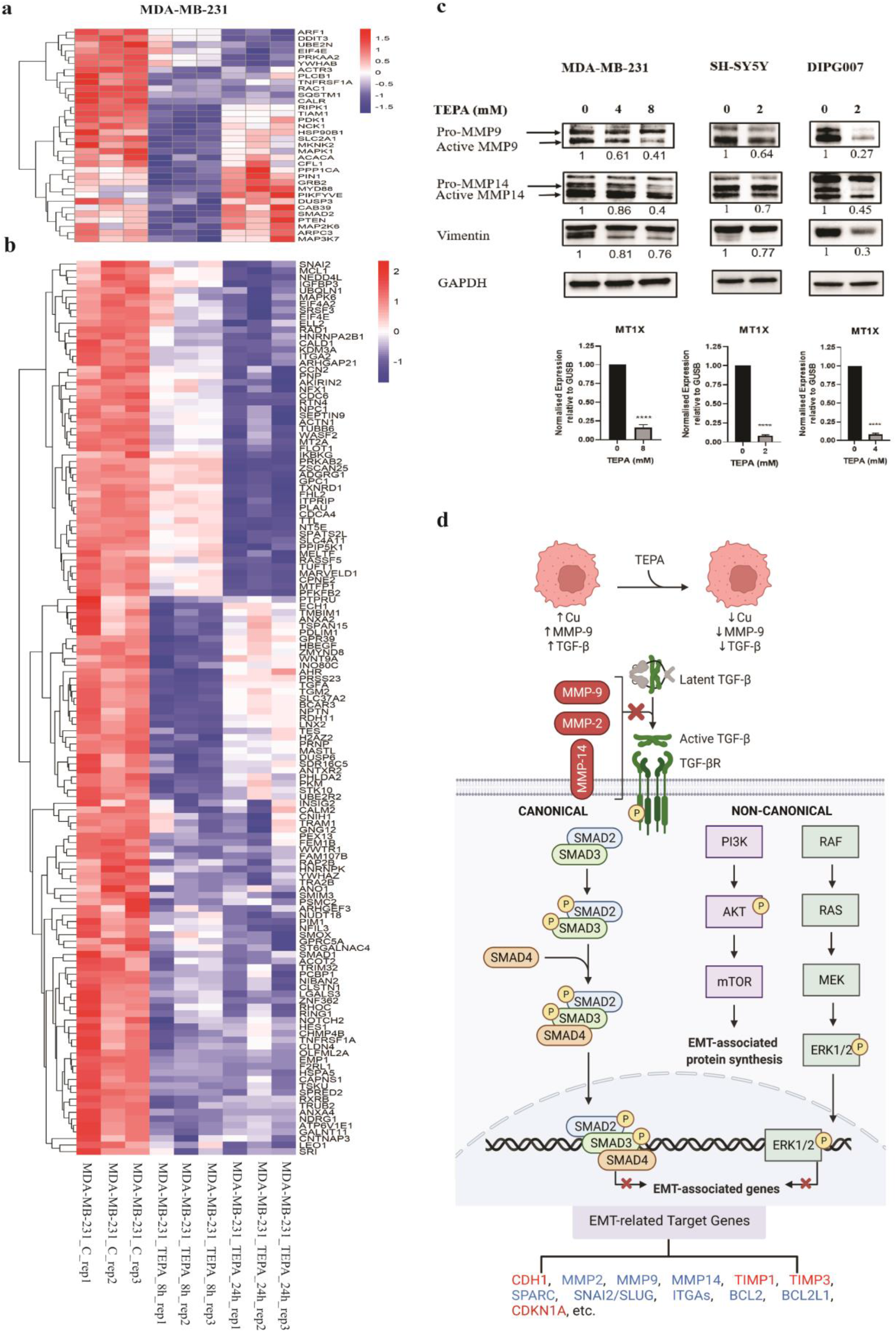
TEPA regulates canonical and non-canonical TGF-β signalling pathways to inhibit EMT and suppress tumorigenesis in MDA-MB-231 cells. Heatmap highlighting TGF-β/SMAD, TGF-β/AKT/mTOR and TGF-β/RAS/RAF/MEK/ERK signalling pathways’ genes deregulation using RNAseq data analysis for MDA-MB-231 cells treated with TEPA for 8 hours (a) and 24 hours (b) compared to the non-treated controls. Heatmap colour scale is proportional to row-scaled VST-normalized expression. RNAseq data revealed that many target genes related to reversing EMT, immune evasion, and suppressing tumor growth downstream of TGF-β /SMAD, TGF-β/AKT/mTOR and TGF-β/RAS/RAF/MEK/ERK signalling are dysregulated following treating cells with TEPA. c Analysis of MMP9, MMP14, and vimentin at protein levels. Western blot analysis of MMP9, MMP14, and vimentin following treatment of MDA-MB-231, and SH-SY5Y, DIPG007, and DIPG010 cells with TEPA (0, 4 & 8mM for MDA-MB-231 and DIPG010, and 0 & 2mM for SH-SY5Y and DIPG007) for 24 hours. GAPDH has been used as a positive and an endogenous control. Down, Expression analysis by quantitative PCR. The alteration of MT1X gene expression following treating MDA-MB-231, SH-SY5Y, and DIPG007 cells with TEPA for 24 hours were analyzed by Real-Time PCR. MT1X is a surrogate marker which its expression is proportional to the intracellular copper concentration. GUSB was used as an internal control. d Schematic of tumor suppression mechanism by TEPA. TEPA has a regulatory effect on tumor suppression by attenuating the canonical TGF-B/pSMAD2/3 and non-canonical TGF-β/AKT/mTOR and TGF-β/RAS/RAF/MEK/ERK signalling pathways and downstream target genes related to cancer initiation and progression and immune evasion. Based on previous studies, MMPs are responsible for cleaving the inactive form of TGF-β and releasing the active form of TGF-β. Our RNA-seq data revealed that MMP2, MMP9 and MMP14 downregulate and MMPs inhibitors of TIMP1 and TIMP3 upregulate following treating cells with TEPA.

To further validate that TEPA has a robust impact on EMT pathway in cancer cells, we performed western blot analyses. We investigated the activity of TEPA on canonical TGF-β/SMAD2&3 and non-canonical TGF-β/PI3K/AKT and TGF-β/RAS/RAF/MEK/ERK signaling pathways. We found that TEPA downregulated the canonical TGF-β signaling pathway by reduction of both total SMAD2 expression and phosphorylation at the C-terminal of SMAD2 (Fig.4c). Furthermore, we demonstrated the inhibitory effect of TEPA on the TGF-β non-canonical pathway by decreasing AKT and ERK phosphorylation (Fig.4c). This inhibitory effect of TEPA on TGF-β is also evidenced by a decrease in vimentin, and MMP9 and MMP14, which play a key role in promoting tumor invasion in cell lines (Fig. 5c). Moreover, we found a downregulation of MMP2, mTOR and phospho-mTOR proteins and an upregulation of E-cadherin in MDA-MB-231 cells after TEPA treatment (Suppl Fig. 4). Finally, to demonstrate the activity of TEPA we showed a reduced expression of MT1X, a metallothionine used as surrogate measure of levels of copper, by real-time PCR (Fig.5c). Taken together, our data confirmed that copper chelation with TEPA significantly reduces EMT-related metastasis via attenuating TGF-β /pSmad2, and TGF-β /PI3K/pAKT and TGF-β/RAS/RAF/MEK/ERK pathways in NB, DIPG and TNBC cancer cells.

### TEPA inhibits the *in vivo* tumor growth in 4T1.2 syngeneic murine model of triple-negative breast cancer

We evaluated the anti-cancer activity of TEPA using a fully immunocompetent murine pre-clinical syngeneic model of TNBC. 4T1.2 is a highly aggressive murine TNBC that recapitulates human TNBC tumor phenotypes with high propensity for spontaneous metastasis. 4T1.2 cells were implanted into the 4th mammary fat pad of 6-weeks old female Balb/c mice, and once tumor size reached approximately 50 mm^3^, mice were treated with either vehicle (saline) or TEPA (400 mg/kg and 800 mg/kg) daily by oral gavage. We used same the same concentrations of TEPA already tested in the neuroblastoma mouse model (38). We saw no impact of TEPA (400 mg/kg) on primary tumor growth, whereas 800 mg/kg TEPA significantly inhibited the 4T1.2 tumor growth compared to vehicle-treated control mice (Fig. 6a). TEPA was well-tolerated in mice as evidenced by no significant change in body weight.

**Figure 6.**
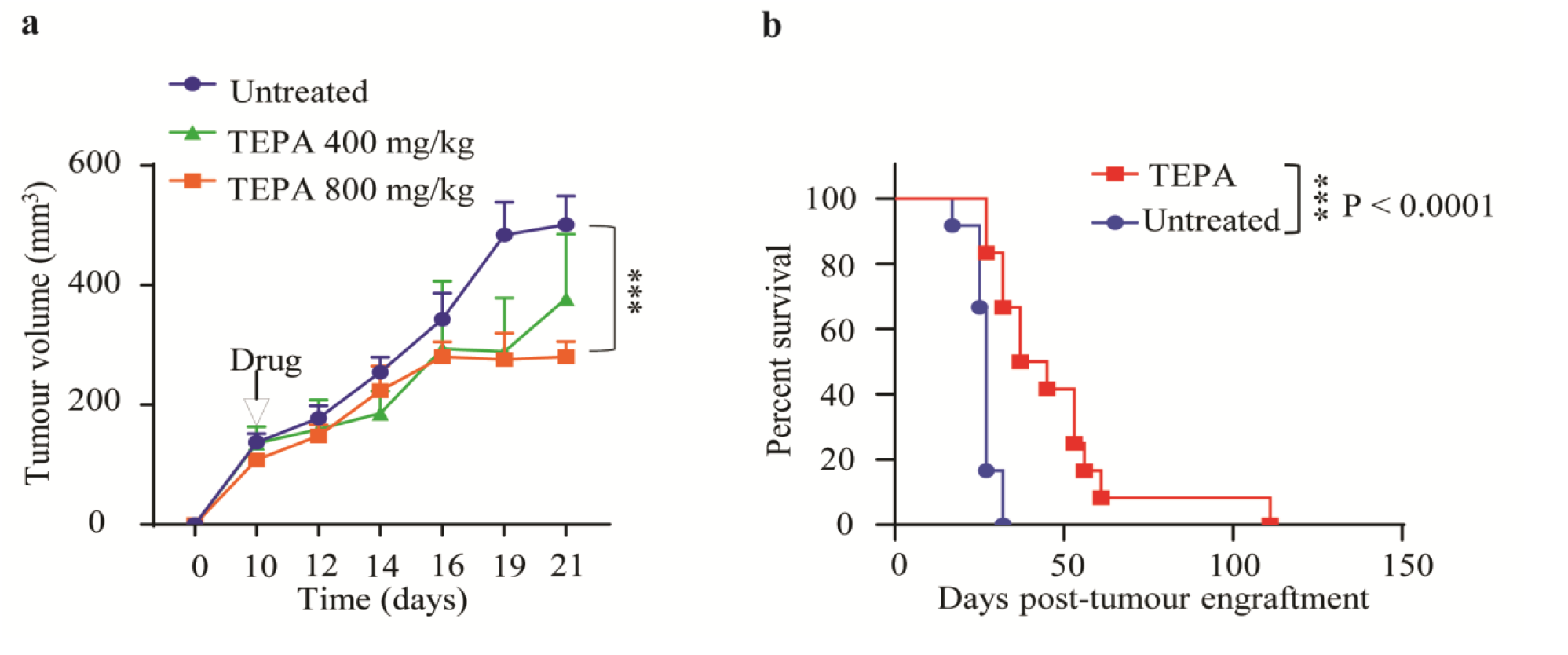
TEPA inhibits the in vivo tumor growth in syngeneic model of triple-negative breast cancer. a Murine 4T1.2 syngeneic TNBC tumor growth following treatment with vehicle or TEPA (400 mg/kg and 800 mg/kg, daily, oral gavage) for three weeks is shown. Treatment was started once tumors became palpable (50-100 mm3). The mean tumor volume of each treatment group from each mouse is presented (n=6 mice/group). Two-way ANOVA was followed by Sidak’s post-test for tumor growth analysis. b The survival analysis for murine 4T1.2 syngeneic TNBC tumors following treatment with vehicle or TEPA (800 mg/kg, daily, oral gavage) for three weeks is calculated by the Kaplan-Meier survival analysis. Mice were treated with TEPA for three weeks, treatment was withdrawn, and mice were followed for survival until tumors reached 1000 mm3.

We next examined the effect of TEPA treatment on the overall survival of 4T1.2 tumor-bearing mice. 4T1.2 tumor-bearing mice were either treated with vehicle or 800 mg/kg TEPA. After 21 days, we withdrew the treatment and followed the mice for survival. We found that 800 mg/kg TEPA, significantly prolonged the survival in the 4T1.2 tumor-bearing mice (Fig. 6b). The median survival time in the control mice was 27 days, where TEPA increased median survival to 41 days (p – 0.0002). Moreover, this experiment showed us that TEPA at concentration of 400mg/kg did not reduced the primary tumor, suggesting this as right dose to use to study a potential effect on EMT and metastasis *in vivo*.

### TEPA reduces lung metastases in 4T1BR metastatic TNBC model *in vivo*

Based on our *in vitro* findings that TEPA inhibits EMT and significantly inhibits the migration of highly metastatic cancer cells, we aimed to investigate the effect of TEPA on inhibiting metastases *in vivo*. We investigated the effect of TEPA on lung metastasis using an aggressive syngeneic murine 4T1BR4 model for TNBC. The 4T1BR4 cells are derived from the parental 4T1 cells however with a higher engraftment rate in the lungs(67). We used TEPA at the reduced 400 mg/kg as we had observed this concentration not to have an effect in reducing the tumor growth of the primary tumor (Fig.7a). We found that lung metastasis was significantly reduced in lung nodules in mice treated with TEPA compared to their vehicle treated controls (P value>0.001 Fig.7b & 7c). Our data convincingly show that TEPA exerts potent cytotoxicity on metastatic TNBCs, reducing the overall lung metastatic burden (Fig.7b & 7c). In addition, a significant reduction was observed in TGF-β levels in the serum of the mice in response to TEPA treatment (Fig. 7d). This is also consistent with analysis performed in the NB TH-MYCN mouse model, where we observed decreased levels of TGF-β after treatment with TEPA (Suppl Fig. 5). This is likely a consequence of the effect of TEPA in inhibiting TGF-β signaling.

**Figure 7.**
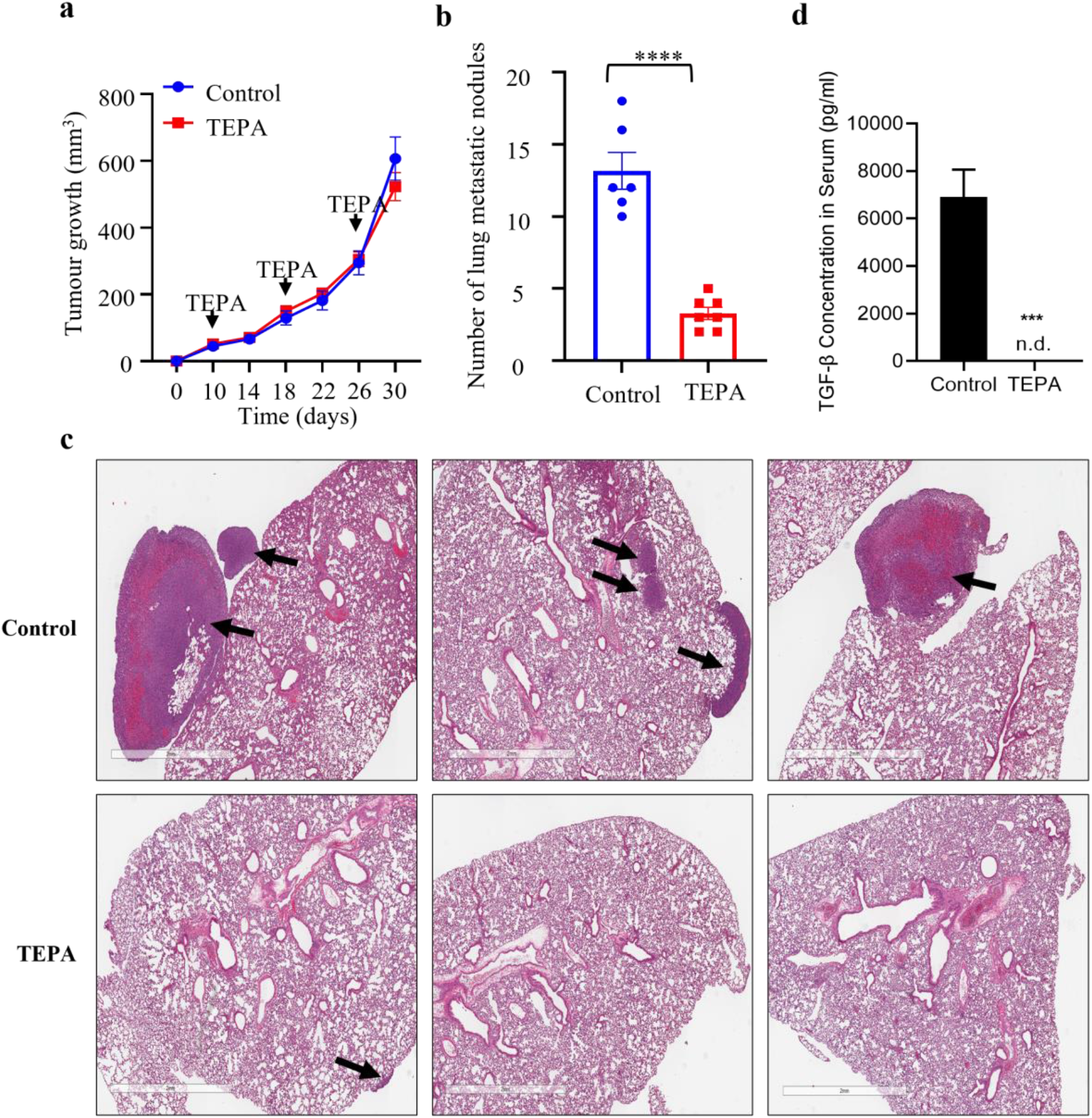
TEPA reduces lung metastases in 4T1BR metastatic TNBC model in vivo. Murine 4T1BR tumor-bearing mice were treated with vehicle or TEPA (400 mg/kg, daily, oral gavage) for three weeks. At the end of the treatment, lungs were isolated and stained with H&E to analyse tumor metastases. a Tumor growth analysis of murine 4T1BR tumor-bearing mice treated with vehicle or TEPA (400 mg/kg). Two-way ANOVA was followed by Sidak’s post-test for tumor growth analysis (n=6). b Quantification of the number of metastatic lung nodules in the vehicle and TEPA-treated mice. Unpaired “t” test was performed (n=6). c Representative images of lung metastasis in 4T1BR tumor model treated with the vehicle or TEPA. The metastatic nodules were stained with H&E. d The protein level of TGF-*β* in the sera of 4T1BR syngeneic mice treated with TEPA (400 mg/Kg, daily, oral gavage) for 30 days.

## Discussion

Copper is a crucial metal acting as a co-factor for many metalloenzymes and proteins involved in several cancer de-regulated pathways e.g. angiogenesis and red-ox status, promoting tumor progression in a large set of tumors like breast, lung, colon and melanoma cancers. Copper chelation has efficacy in several cancer types in clinical trials and recently copper chelation therapy has been proposed as a potential adjuvant strategy to overcome BRAF-MEK1/2 inhibition gained resistance in melanoma and colon cancer (68, 69). We previously reported that the copper chelator TEPA was able to reduce tumor growth and PD-L1 expression increasing tumor-infiltrating immune cells, in a neuroblastoma mouse model (70). As copper chelators have multiple mechanisms of action, we aimed to determine mechanisms which would contribute to explaining copper chelation’s anti-cancer impact across different cancer types. Using transcriptomic data from three different tumor types and additional supporting studies, we determined EMT and TGF-β signaling to be a previously undescribed target of copper chelation in cancer.

EMT is a reversible process in which epithelial cells lose their cellular properties towards a more mesenchymal phenotype, allowing invasion, cell migration, tumor development, and metastasis (71) (72). TGF-β exerts its effect on promoting EMT via canonical (SMADs) and non-canonical (PI3K/AKT/mTOR & RAF/RAS/MEK/ERK) TGF-β signaling pathways. In this study, we have demonstrated that copper chelation can inhibit epithelial-mesenchymal transition and shift cell properties from a mesenchymal state to a more epithelial-like phenotype in TNBC, NB and DIPG cancer cells via attenuating TGF-β signaling pathway activation and impair tumor invasiveness. Diminished intracellular copper availability influenced both canonical and non-canonical TGF-β signaling pathways perturbing the expression of several key genes involved in EMT such as TGF-β1, MMP2, MMP9, MMP14, E-cadherin, vimentin, SPARC, ZYX, VCAN, ITGAs, as well as a crucial transcription factor responsible for the mesenchymal fate (SNAI2/SLUG). Interestingly TEPA treatment also changed the location of EMT markers SNAI1 and ZEB1 which are crucial transcription factors in EMT process. Thus a shift of location proportions in favor of the cytoplasm would suggest their gene transactivation activity on DNA is reduced. These changes of EMT markers were functionally validated by our wound healing and migration assay results. Both TNBC and NB cell lines displayed decreased migration with TEPA treatment.

TGF-β is usually secreted into the extracellular matrix as an inactive form (latent TGF-β) where MMP9 and MMP14 cleave latent TGF-β and release the active form of TGF-β in the extracellular matrix triggering EMT tumor (73-76) (37, 77). Copper chelation with TEPA decreased both MMP9 and MMP14 active forms in MDA-MB-231, SH-SY5Y, and DIPG007 and MMP2 in MDA-MB-231 and DIPG007 cells, exerting an alternative level of control on TGF-β function in addition to TEPA direct attenuation of TGF-β transcription. Additionally, TEPA significantly increased the expression of TIMP3 and TIMP4 in TNBC, and TIMP1 in NB cells 24 hours after treatment. TIMP1, TIMP3, and TIMP4 are important inhibitors of MMP-9 protein (78, 79). TIMP proteins bind to the active site of MMPs and inhibit their activity by blocking binding of MMPs to ECM substrates (80).

Lowering copper also induced downregulation of a broad panel of genes whose products impact many critical cancer-related signaling pathways, as depicted in Table 1, including apoptosis, angiogenesis, and inflammation by reducing BCL2 and VEGF protein expression, respectively. A list of the genes and proteins’ expression affected by TEPA in four studied cell lines has depicted in the Table 2.

**Table 2.** List of genes and proteins’ phosphorylation affected by TEPA in RNA-seq, Real-Time PCR, and western blot analysis of MDA-MB-231, SH-SY5Y, DIPG007, and DIPG010 cell lines. These genes and proteins are important in the migration, and invasion of cancer cells via dysregulation of TGF-β and EMT pathways.

| Cell lines | Genes / Protein Symbol |  |
| --- | --- | --- |
|  | Downregulated | Upregulated |
| MDA-MB-231 | TGF-β1, TGFBR2, SPARC, SNAI2, SMAD2, p-SMAD2 MMP2, MMP9, MMP14, MMP11, MMP15, Vimentin, RAC2, BCL2, PLOD2, PLAT, MARCKSL1, CCR7, PAK1, ZYX, LAMA4, p-AKT, p-ERK1/2 | CDKN1A, TIMP3, TIMP4, E-cadherin (CDH1) |
| SH-SY5Y | TGF-β1, p-SMAD2, SMAD2, MMP9, MMP14, Vimentin, ZEB1, ZYX, RAC1, PLAT, p-AKT, p-ERK1/2 | TGIF1, CDKN1A |
| DIPG007 | SMAD2, p-SMAD2, MMP2, MMP9, MMP14, Vimentin, ZYX, VCAN, PXN, AKT, p-AKT, ERK1/2, p-ERK1/2 | CDKN1C |
| DIPG010 | TGF-β1, SPARC, N-cadherin (CDH2), Vimentin, MMP2, ZYX, LAMA4, PLAT, VCAN | |

We also decided to test TEPA in the 4T1BR4 TNBC mouse model since this is a well-known copper-dependent cancer and other copper chelators have been used successfully in a clinical trial in TNBC patients (81). We observed that at a concentration of the drug, which did not have an effect on reducing the primary tumor was remarkably able to reduce metastasis. When we analyzed the cytokine profile and RNA-seq data from NB, TNBC and DIPG cell lines, TGF-β was significantly reduced in all cancer types as were the EMT-related pathways which were significantly down-regulated.

It is interesting that TGF-β is reported to play an important role in cancer immune evasion (82). TGF-β suppresses the immune system by increasing PDL1 expression, which is dependent mainly on the activation of PI3K/AKT pathway (83). This action is consistent with our previous finding in neuroblastoma that copper chelation altered tumor immune evasion and reduced PDL1 expression (70). Transcriptomic data from TEPA treated breast cancer cells also displays changes in the immune-related genes (including IL6, IL6R, IL15, CXCL1, CXCL8, and PTX3) in breast cancer. Future studies are needed to ascertain if the copper chelation inhibition of TGF-βsignaling is required to alter the immune evasion previously reported in the neuroblastoma context or even as a general anti-cancer mechanism. This aspect may have led to improved immune surveillance and subsequently contributed to inhibition of metastatic spread in our *in vivo* model and may indicate that copper chelation could be used in place of, or in combination with immunomodulation cancer therapy regimes.

21 drugs targeting TGF-β have failed in clinical trials, partly because of severe side effects arising in patients (44-48). This raises concerns over the translational feasibility of the current bench-to-bedside strategies directly targeting TGF-β. Here, the copper chelator TEPA, analogue of the FDA-approved Trientine, significantly reduces TGF-β levels and downstream signaling pathways. This presents copper chelation as a non-toxic and easily delivered alternative to direct targeting of TGF-β.

In conclusion, we report for the first time, that the copper chelator TEPA causes depletion of TGF-β, which results in inhibition of metastasis. TEPA influences EMT *in vitro*, in three tumor types, and more remarkable, low doses of TEPA, despite not impacting tumor growth, completely block TNBC lung metastasis. TGF-β signaling is a drug target in most cancer types and several TGF-β targeting agents have shown promise in pre-clinical models, but the results in clinical trials are disappointing because of their strong side effects (84). Considering that copper chelating agents are already used for Wilson's disease, are non-toxic and can deplete copper from all tissues of the body, this study provides an attractive framework for a new therapeutic strategy to target TGF-β and subsequently impact both tumor growth and metastatic potential.

## Acknowledgements

The authors acknowledge financial support from: National Health and Medical Research Council (NHMRC) Career Development Fellowship (APP1164960); Cure Cancer Institute NSW Career Development Fellowship (RG161875); Cure Cancer Australia grant (RG180859)

