## Supplementary Figure 1 for "Copper chelation inhibits TGF-*β* pathways and suppresses epithelial-mesenchymal transition in cancer"

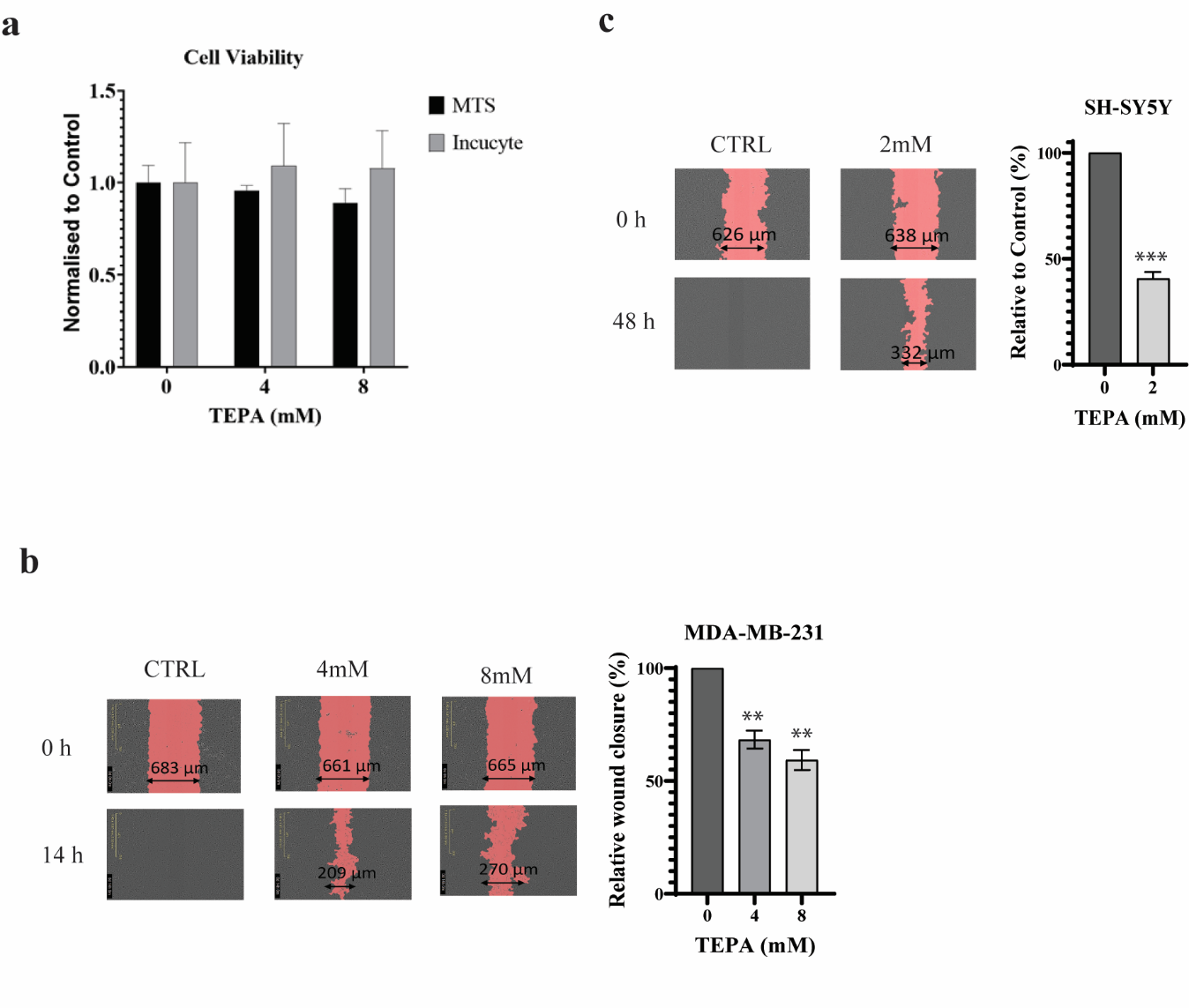

**Suppl Figure1**- Analysis of cell viability and migration following treating cells with TEPA. **a** MTS assay was performed on MDA-MB-231 cells treated with 0, 4, and 8 mM TEPA for 24 hours. The absorbance assay was read at 490nm using the Benchmark Plus Microplate Spectrophotometer System. All results were normalized to the control with error bars representing the standard error between 2 independent experiments with three technical replicates per experiment. **b & c** Cell migration analysis by scratch wound assay for MDA-MB-231 and SH-SY5Y cells, respectively. Pink areas indicate the scratch-wound area, and gray areas show the cell monolayer. Cell migration into the wound area is represented by wound width (μm) in the scratch-wound assay images. Graphs represent the percentage of wound closure observed over the corresponding time points. Results were normalized to the 0h time point as the control. Significance was determined by one-way ANOVA with p-value <0.0001 for MDA-MB-231 cells and Paired t-test with p-value= 0.0009 for SH-SY5Y.
