## Supplementary Figure 2 for "Copper chelation inhibits TGF-*β* pathways and suppresses epithelial-mesenchymal transition in cancer"

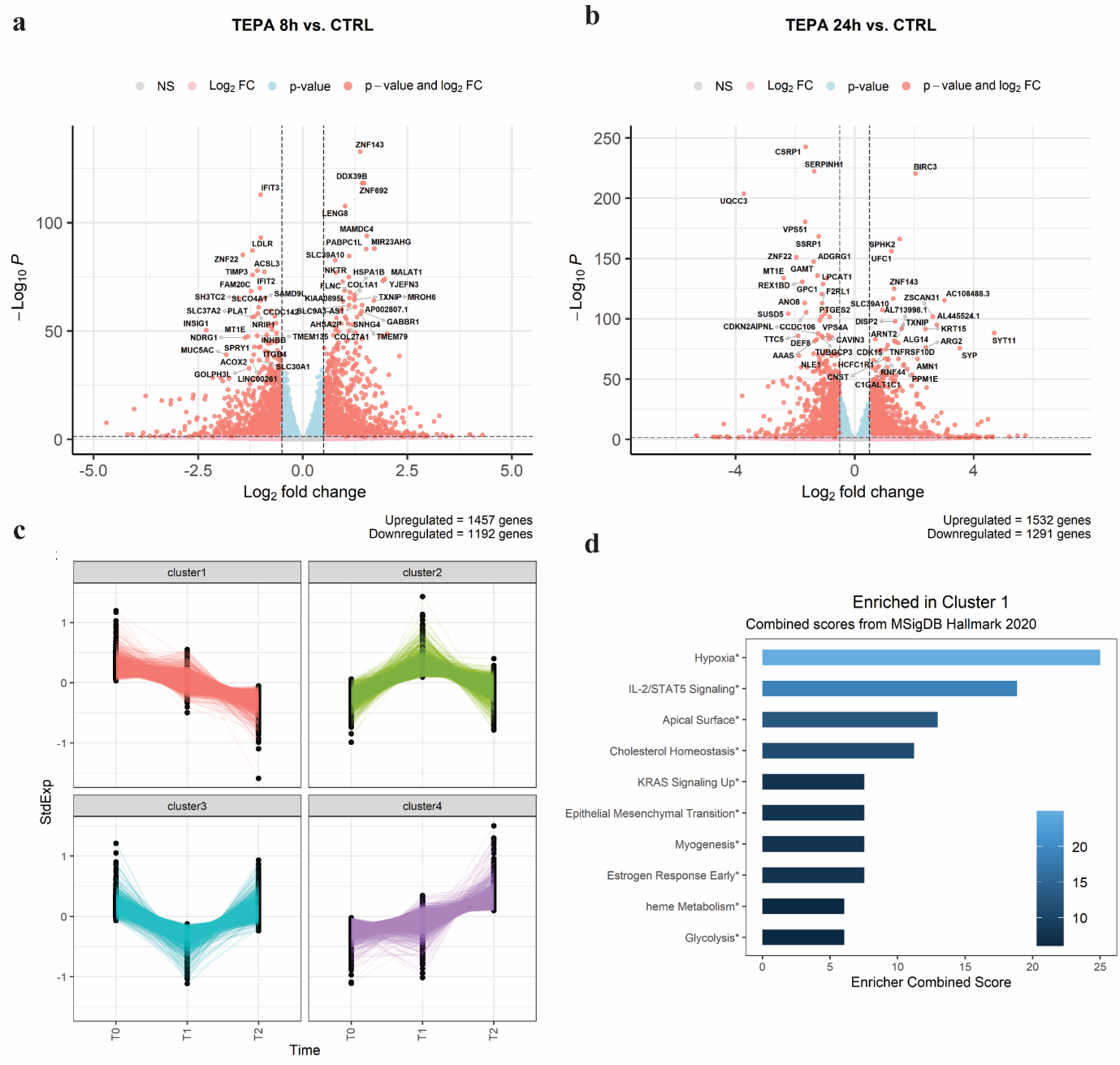

**Suppl Figure 2-** Gene expression changes in TEPA-treated MDA-MB-231 cells. Volcano plots showing gene expression changes after 8 (**a**) and 24 (**b**) hours of TEPA treatment. The top significant 25 up and top 25 down-regulated genes are labeled. **c** K-means clustering classifies gene expression changes into 4 clusters. Genes included in cluster 4 show a time-dependent downregulation. **d** Top 10 MSigDB Hallmark enriched gene sets. Significant enrichments of MSigDB gene sets were evaluated through enrichment analysis performed with EnrichR. Color intensity is referred to the enrichment score computed by EnrichR and calculated as follows: combined score = log(p) * z, where p is the Fisher exact test p-value, and z is the z-score for deviation from expected rank. Asterisks mean a corrected p-value for multiple testing < 0.05.
