## Supplementary Figure 3 for "Copper chelation inhibits TGF-*β* pathways and suppresses epithelial-mesenchymal transition in cancer"

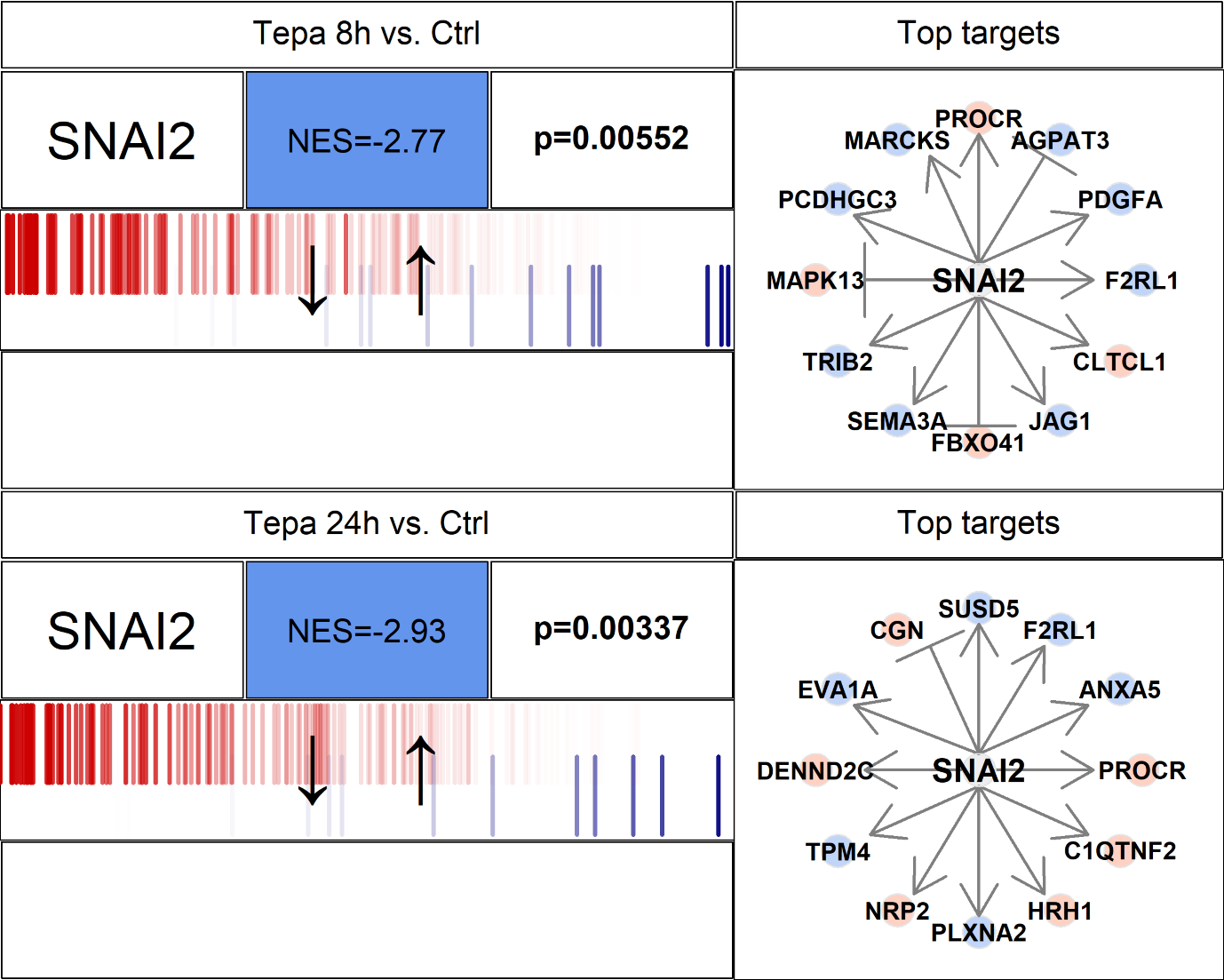

**Suppl. Figure 3- Master Regulator Analysis.** SNAI2 sub-network downregulation in TEPA-treated cells at 8 (upper panel) and 24 hours (lower panel). The top 12 highest-likelihood targets are shown on the right side. The genes in each network are shown in a barcode-like diagram showing all transcriptome genes by means of their differential expression upon TEPA treatment, from the most downregulated (left) to the most upregulated(right). A blue background on the NES box is used to indicate a negative enrichment (or repression of the corresponding co-expression network).
