## Supplementary Figure 4 for "Copper chelation inhibits TGF-*β* pathways and suppresses epithelial-mesenchymal transition in cancer"

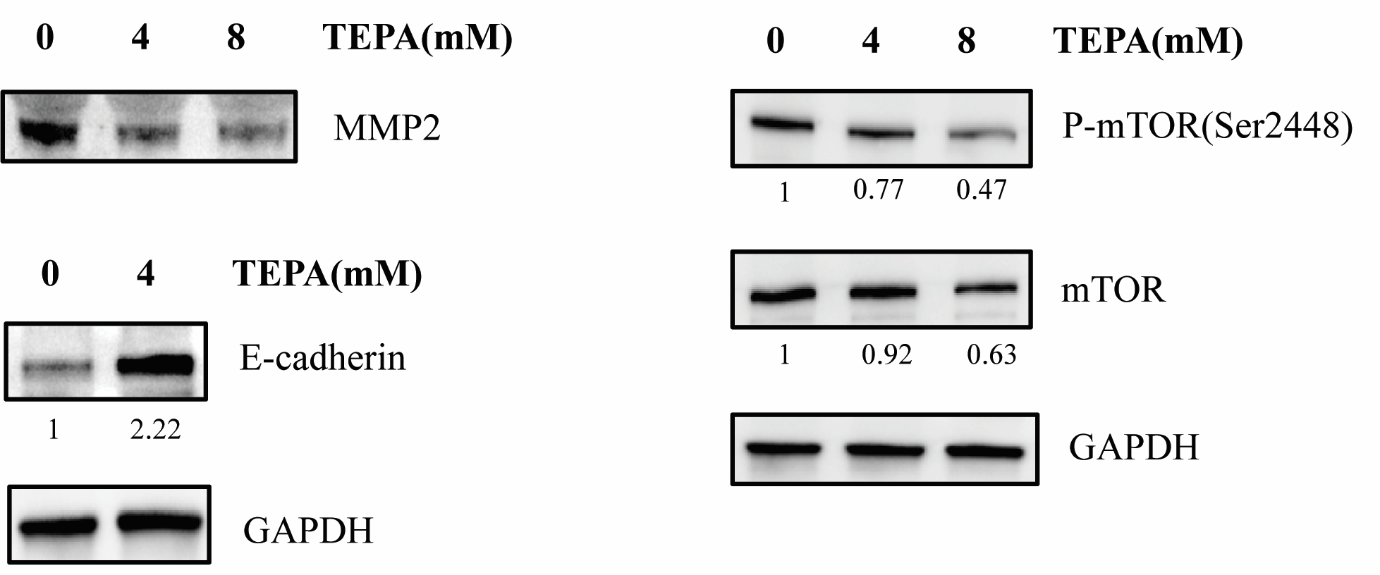

### **Suppl. Figure 4-** Western blot analysis of MMP2 in MDA-MB-231 cell supernatant, and mTOR, phospho-mTOR (Ser2448), and E-cadherin (CDH1) in cell lysate. For MMP2, MDA-MB-231 cells were treated with TEPA (0, 4, and 8mM) in serum-free media for 24 hours. Then cell supernatants were collected, and soluble proteins were concentrated using Ultracel-10 regenerated cellulose membrane (Amicon® Ultra-15 Centrifugal Filter Unit, Merck). 20 µg of each sample was used for western blot analysis with indicated antibodies.
