## Supplementary Figure 5 for "Copper chelation inhibits TGF-*β* pathways and suppresses epithelial-mesenchymal transition in cancer"

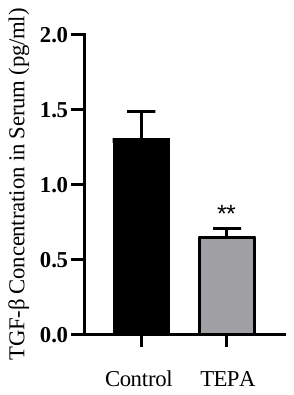

**Suppl Figure 5.** The expression level of TGF-β in the TH-MYCN neuroblastoma mouse model treated with TEPA. TH-MYCN mice were treated with 400 mg/Kg TEPA for 7 days and the TGF-β expression level was analyzed in the sera of mice using a multiplex cytokine assay. Significance was determined by unpaired t-test with p-value=0.0099.
